# Quantitative analysis of tyrosine phosphorylation from FFPE tissues reveals patient specific signaling networks

**DOI:** 10.1101/2020.09.10.291922

**Authors:** Ishwar N. Kohale, Danielle M. Burgenske, Ann C. Mladek, Katrina K. Bakken, Jenevieve Kuang, Judy C. Boughey, Liewei Wang, Jodi M. Carter, Eric B. Haura, Matthew P. Goetz, Jann N. Sarkaria, Forest M. White

## Abstract

Formalin fixed paraffin embedded (FFPE) tissues are an invaluable source of clinical specimens. Tyrosine phosphorylation (pTyr) plays a fundamental role in cellular processes and is commonly dysregulated in cancer but has not been studied to date in FFPE samples. We describe a method for quantitative analysis of pTyr signaling networks at an unprecedented sensitivity, with hundreds of sites quantified from 1-2 10-μm sections of FFPE tissue specimens. Phosphotyrosine profiles of flash frozen and FFPE tissues derived from the same tumors suggest that FFPE tissues preserve pTyr signaling characteristics in PDX tumors and archived clinical specimens. Differential activation of oncogenic proteins was observed in triple negative breast cancer tumors as well as lung cancer tumors, highlighting patient specific oncogenic driving kinases and indicating potential targeted therapies for each patient. These data highlight the capability for direct translational insight from pTyr analysis of small amounts of FFPE tumor tissue specimens.

## Introduction

The rise of targeted therapeutics over the past two decades has highlighted a need for personalized cancer medicine to match optimal therapy to each patient. Precision medicine has commonly relied on genomic and transcriptomic tumor profiling^1,2^, yet these approaches have yielded limited success, possibly due to incomplete systems biology characterization of the tumors. While these “omics” approaches provide information on genomic mutations or altered transcript expression, neither approach directly measures signaling networks that drive tumor progression and regulate inherent and acquired therapeutic resistance. Analysis of phosphorylation mediated signaling networks can provide crucial information on oncogenic drivers or dysregulated networks in patients^3^.

Tyrosine phosphorylation (pTyr) accounts for only 0.1-1% of the total phosphoproteome, is highly conserved and tightly regulated, and controls many aspects of cellular and tumor biology^4,5^. Thirty percent of the known oncoproteins are tyrosine kinases (TK)^6^, and their disproportionate role in oncology has led to development of many TK inhibitors (TKIs)^7^. Quantitation of tyrosine phosphorylation can measure activated signaling networks in patient tumors and therefore highlight particular therapeutic options. However, such analysis has historically been limited by the large amounts of clinical tissue required, and the need for frozen tissues, both of which can be challenging to obtain routinely.

Human tissue specimens obtained from diagnostic and surgical procedures are commonly preserved as formalin fixed paraffin embedded (FFPE) tissues in the clinic, and FFPE tissues are readily available in tumor tissue banks^8^. While protein expression and global phosphorylation are increasingly being studied in FFPE tissues^9–12^, global phosphorylation enrichment techniques typically yield few tyrosine phosphorylation sites due to its low abundance. Additionally, given previous reports of post-surgical ischemic effects on phosphorylation^13,14^, it is not known how well tyrosine phosphorylation is preserved in FFPE tissues. Therefore, a comprehensive comparison of FFPE and flash frozen tissue is required to determine whether FFPE tissues can provide accurate quantification of cell signaling networks.

We have developed an approach enabling tyrosine phosphorylation profiling of FFPE tissues with unprecedented sensitivity. We demonstrate quantification of ∼2000 pTyr sites belonging to critical cancer pathways from multiple 10-μm sections of FFPE tissues, representing a ∼20-fold improvement in sensitivity for pTyr analysis. To understand the effects of FFPE preservation on pTyr levels, we compared pTyr profiles of flash frozen and FFPE tissues derived from the same tumors and show that FFPE tissues faithfully preserve most, but not all, pTyr signaling. Using our optimized protocol, >900 pTyr sites were quantified from single tissue punches obtained from clinical FFPE blocks containing triple negative breast cancers (TNBC) specimens, and from 1 or 2 10-μm sections of FFPE from non-small cell lung cancer (NSCLC) patient tissues. Differential activation of oncogenic proteins such as EGFR, SRC and MET was observed in different TNBC tumors, highlighting putative patient specific oncogenic driving kinases for consideration of targeted therapeutic approaches. These results highlight the direct translational potential of pTyr analysis of FFPE tumor tissue specimens.

## Results

### Phosphotyrosine analysis of FFPE samples is feasible and provides quantitative data on underlying biology

We set out to develop a method for the quantitative characterization of pTyr signaling networks from small amounts of FFPE tissues, and to determine whether the signaling networks quantified from these tissue specimens could provide relevant biological insights. To this end, we developed a protocol combining 2,2,2-trifluoroethanol (TFE) based protein extraction^11^ with paramagnetic SP3 beads based sample processing^9,10^ that allows for robust and sensitive phosphoproteomic analysis of FFPE tissues (Figure 1a). This optimized protocol led to peptide yields of approximately 2 μg per mm^2^ of a 10-μm section of FFPE tissue, and scales with larger tissue sections of FFPE blocks (Adj R^2^ = 0.95, Figure S1a). To assess the feasibility of quantifying pTyr signaling in FFPE tissues, we enriched pTyr-containing peptides through a 2-step protocol, using anti-pTyr antibodies for immunoprecipitation (IP) and immobilized metal affinity chromatography (IMAC) to remove non-specifically retained non-phosphorylated peptides prior to analysis by liquid chromatography tandem mass spectrometry (LC-MS/MS). Using this platform, we performed MS analysis of enriched pTyr peptides from multiple 10-μm sections from patient derived xenograft (PDX) glioblastoma (GBM) tumors (GBM6)^15^. This approach led to identification of 1009, 1031, 1704 and 2165 pTyr sites in 4 different tumors (Figure 1b, Table S1-4), including pTyr sites on 129 kinases spanning across multiple branches of the kinome tree (Figure 1c). Not surprisingly, identified pTyr-proteins provided good coverage of receptor tyrosine kinases including EGFR, an oncogenic driver kinase in GMB6 because of EGFRvIII amplification^15^. Gene ontology analysis of pTyr-proteins indicated enrichment of multiple reactome pathways including axon guidance, signaling by Rho GTPases, signaling by RTKs, cell cycle and cellular senescence, providing further evidence that pTyr enrichment and analysis yields insight into activated cellular pathways in tumors (Figure 1d).

**Figure 1.**
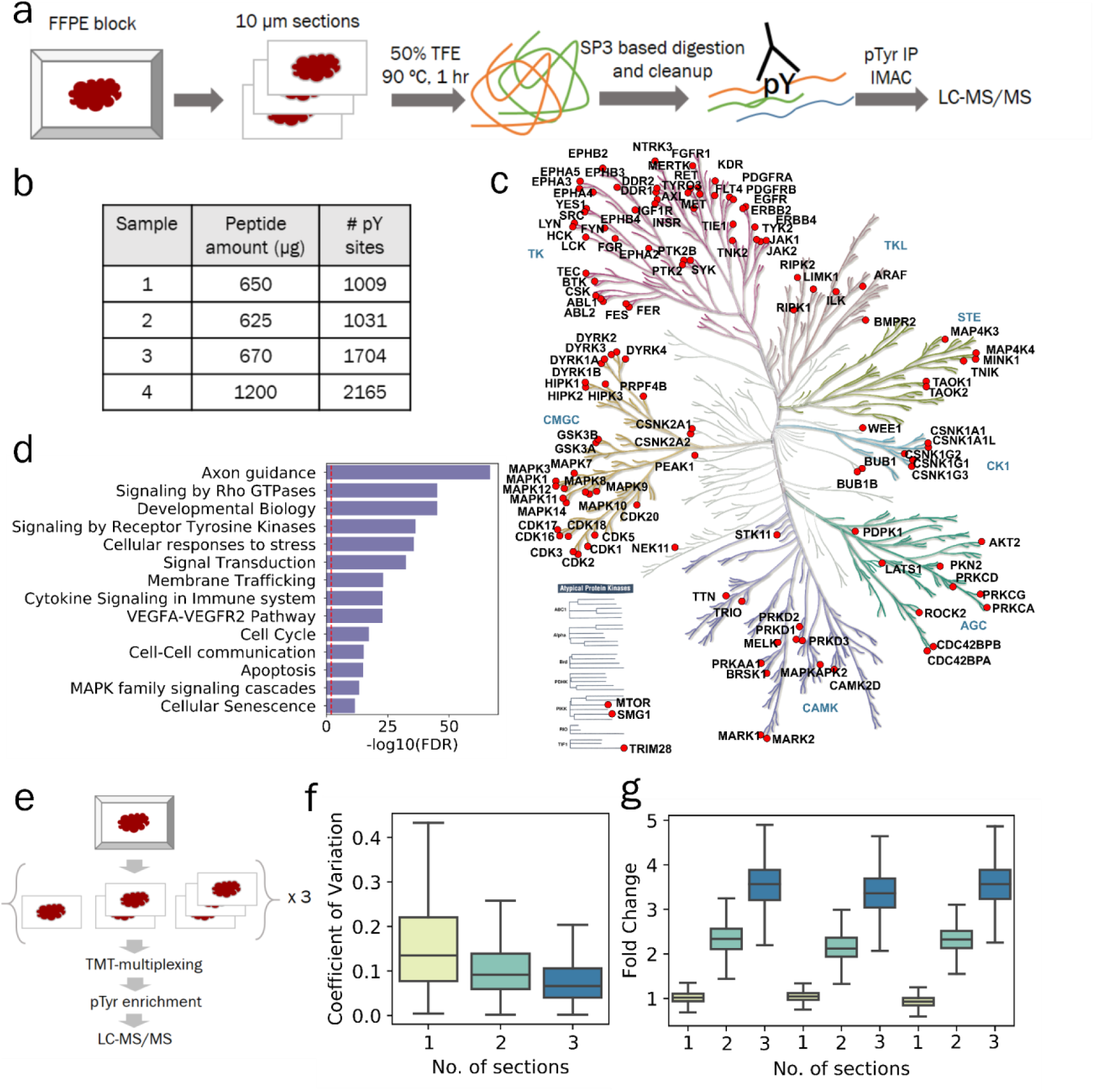
Phosphotyrosine analysis from 10-μm sections of FFPE tissues. **a)** Optimized workflow for extraction and digestion of proteins from FFPE tissues followed by 2-step enrichment of pTyr peptides for LC-MS/MS analysis. **b)** Number of pTyr sites identified from multiple 10-μm sections of PDX tumors. **c)** Kinome tree depicting pTyr containing proteins identified in PDX tumors. **d)** Selected reactome pathways enriched in gene ontology analysis of pTyr-proteins. Dashed red line depicts FDR = 0.01. **e)** Schematic workflow for a multiplexed pTyr analysis of 1, 2 or 3 10-μm sections of FFPE tissues in triplicate. **f)** Coefficient of variation observed across multiple sections of FFPE tissues. Median CV for 1, 2 and 3 sections were 13.5%, 9.2% and 6.7%, respectively. **g)** Fold change of TMT intensities of peptides quantified in each channel compared to the average of TMT intensities from single sections on a peptide basis. Error bars represent interquartile range.

Routine pTyr and global phosphoproteomic analysis of clinical specimens has been hampered by the amount of tissue samples required for such experiments. To assess the low-input sample sensitivity and robustness of our FFPE pTyr analysis platform, peptides derived from 1, 2, or 3 10-μm thick sections of FFPE tissue, corresponding to ∼50, 100, or 150 μg of peptides per sample, were labeled with tandem mass tags (TMT) for multiplexing (Figure 1e) prior to pTyr enrichment and LC-MS/MS analysis. This approach led to identification and quantification of 816 pTyr containing peptides across all samples (Table S5). Median coefficient of variation (CV) for peptides quantified across the replicates were 13.5%, 9.2 % and 6.7 % for 1, 2 or 3 sections, respectively, suggesting robust quantitation even with a single 10-μm section (Figure 1f). Median fold change compared to average of single sections for peptides within the same samples were 1.02, 2.32 and 3.57 for 1, 2 or 3 sections (Figure 1g), respectively. Similar relative quantification was also observed for non-phosphopeptides quantified from the supernatant of pTyr IP, suggesting that sample processing of single sections leads to increased sample loss due to the small amount of input material (Figure S1b, Table S6). Quantified phosphoproteins were enriched in RTKs as well as other branches in the kinome tree (Figure S1c), and belong to multiple pathways such as EGFR, FGFR and PI3K signaling, many of which have been already implicated in promoting GBM tumor progression and therapeutic resistance^16–18^ (Figure S1d). Overall, these data suggest that pTyr analysis of a single 10-μm section of FFPE-preserved tissue specimen is feasible and can yield hundreds of pTyr peptides representing a broad swath of GBM tumor biology.

### Comparison of pTyr, pSer/Thr and Protein levels in FFPE and flash frozen tissues

Our initial data indicate that analysis of pTyr signaling in FFPE GBM PDX tumors can yield a large number of identified and quantified pTyr peptides. However, there was some concern that the time required for formalin fixation, which occurs at ∼1 mm/hr^19^, may lead to altered signaling compared to the time required for flash freezing, which occurs on the sub-second/second time scale, especially given previous data suggesting that ischemia alters pTyr signaling within minutes post-resection^13,14^. To understand the effect of FFPE preservation on pTyr signaling, GBM6 PDX tumors were treated *in vivo* with vehicle or afatinib, a second generation EGFR inhibitor, resected, and then half of the tumor was flash frozen in liquid nitrogen, and other half was processed into an FFPE block (Figure 2a). A piece of each flash frozen tumor tissue was homogenized in 8M urea lysis buffer, as per our standard protocol^13,20– 22^. FFPE tissues were lysed in 50% TFE at 90 °C according to the FFPE protocol described above. To control for the effects of the FFPE protein extraction protocol, a separate aliquot of flash frozen tissue was lysed according to the FFPE protocol. Proteins extracted from these three protocols: 1) FFPE, 2) flash frozen lysed in hot TFE (hereafter referred as Frozen-TFE) and 3) flash frozen with standard 8M urea lysis (Frozen-Urea) were digested to peptides, labeled with 16-plex isobaric mass tags (TMTpro), enriched for pTyr, and analyzed by LC-MS/MS. To assess the effect of FFPE preservation on global phosphorylation and protein expression, we quantified phospho-serine/threonine (pSer/Thr) and proteins levels in the same set of samples by fractionating the supernatant from the pTyr IP. Analysis of pTyr signaling led to identification and quantification of 1128, 1085 and 649 peptides in FFPE, Frozen-TFE and Frozen-Urea workflows, respectively (Figure 2b, Table S7-15). Fewer pTyr sites were identified in the Frozen-Urea condition, as only 8 samples were multiplexed together in this analysis as opposed to 16 samples in the FFPE and Frozen-TFE analysis. Peptides derived from the Frozen-Urea workflow had CV of 18% across biological replicates, whereas Frozen-TFE and FFPE workflows had 20% and 30%, respectively, indicating that FFPE preservation may lead to increased variability among samples. We assessed the statistical significance of pTyr-peptides affected by afatinib treatment in each condition and found that 45% of the pTyr-peptides were statistically significantly different between vehicle and afatinib treated groups in the Frozen-Urea condition compared to 26% in Frozen-TFE and 13% in FFPE. Although samples clustered by treatment condition (afatinib vs. vehicle (Figure S2a)), there were some marked differences between FFPE and frozen tissues. Within each treatment condition, frozen pairs (TFE and Urea) derived from the same tumor tended to cluster together, while FFPE counterparts either clustered with each other or clustered with larger sub-clusters. Similarly, correlation between frozen pairs (0.72 ± 0.15) was statistically significantly higher than that of between FFPE and Frozen-TFE (0.46 ± 0.22, P = 0.02) suggesting that pTyr underwent minimal changes during protein extraction steps but more pronounced changes during FFPE preservation (Table S16). These results suggest that some changes in pTyr signaling can occur during FFPE preservation and that these changes can affect our confidence in determining signaling network changes due to drug treatment. In contrast to the pTyr data, pSer/Thr data and protein expression data still clustered by treatment condition, but not by preservation or processing technique (Figure S2b-c), suggesting that these peptides were less affected by sample preservation or processing. Despite similar CV’s, the percentage of peptides that were statistically significantly different in the vehicle vs. afatinib conditions were still lower in FFPE tissues (Figure 2b), possibly because of overall lower TMT intensities for pSer/pThr and unmodified peptides derived from FFPE tissues (Figure S2d-e). Unexpectedly, pTyr-peptide intensities were higher in samples derived from FFPE tissues compared to Frozen-TFE (Figure S2f).

**Figure 2.**
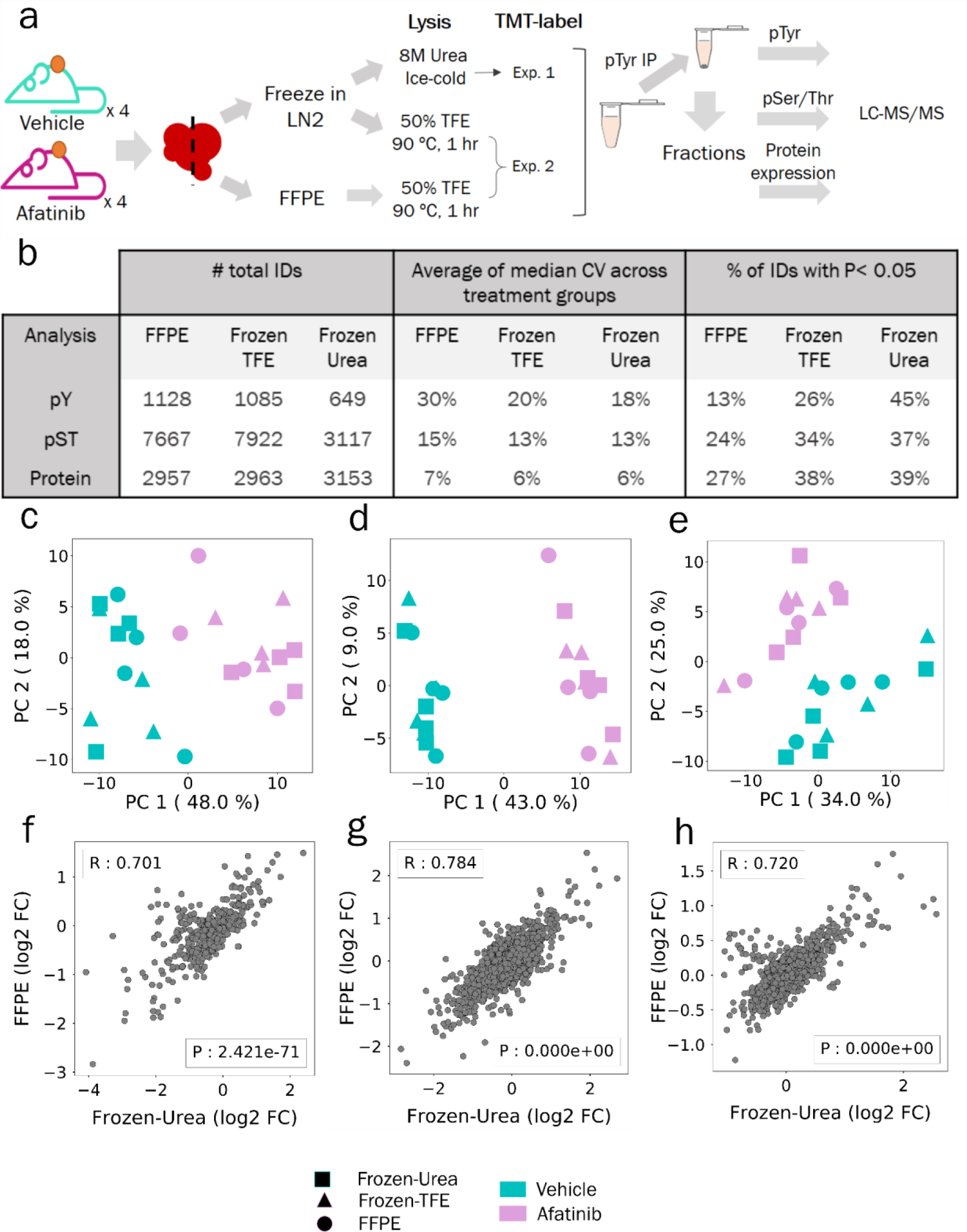
Comparison of pTyr, pSer/Thr and proteins levels in FFPE and flash frozen tissues. **a)** Schematic of experimental design to compare proteomics in FFPE and flash frozen tissues. **b)** Number of unique phospho-peptides or proteins identified and quantified across different workflows with observed CVs and proportion of significantly different IDs between vehicle and afatinib treatments. **c-e)** Principal component analysis of phospho-peptides or proteins quantified across FFPE, Frozen-Urea and Frozen-TFE workflows: **(c)** pTyr (n = 475 peptides), **(d)** pSer/Thr (n = 2283 peptides) and **(e)** protein level (n = 2647 proteins). Quantified levels were mean normalized and log2 transformed within each workflow before concatenating together. **f-h)** Correlation plots of fold changes observed between afatinib and vehicle treated groups in Frozen-Urea samples and their FFPE pairs: **(f)** pTyr, **(g)** pSer/Thr and **(h)** proteins. For each phospho-peptide or protein, fold changes were calculated between mean levels observed in afatinib (n = 4) and vehicle (n = 4) treated groups. R represent Pearson’s correlation.

In order to further assess the effects of sample preservation and processing on signaling and proteomic analysis, we performed principal component analysis (PCA) on peptides or proteins that were quantified across all three workflows. PCA analysis showed that despite different tissue preservation and processing methods pTyr, pSer/Thr and protein levels segregated according to afatinib treatment (Figure 2c-e). We also looked at the correlation between different preservation and processing methods on a peptide-specific basis. Average fold change of treated over the control samples were highly correlated between Frozen-Urea and FFPE tissues with R of 0.70, 0.78 and 0.72 for pTyr, pSer/Thr and protein levels, respectively, underscoring that FFPE tissues preserve similar biology relative to frozen tissues (Figure 2f-h). Taken together, these data suggest that pTyr signaling in FFPE tissues was comparable, but not identical, to the frozen tissues, as some changes in pTyr signaling occur during formalin fixation.

### Similar biology can be read out from FFPE and frozen tissues

Given some differences in pTyr signaling during FFPE preservation, we wanted to understand whether FFPE tissues can still provide similar biological network information compared to frozen tissues. Treatment with afatinib is expected to lead to decreased EGFR activation; accordingly, we detected decreased tyrosine phosphorylation on several proteins in the EGFR pathway including EGFR itself, as well as sites on GAB1, GAB3, SHC1, SHC4, PTPN11 (SHP2), MAPK1 (ERK2) and MAPK3 (ERK1), all of which decreased by 2-10 fold in afatinib treated tumors compared to vehicle control (Figure 3a). Additionally, other proteins directly downstream of EGFR such as CBL (endocytosis), PLCG1 (phospholipase c pathway), PIK3R2 (cell survival) and STAT5A/B (JAK-STAT pathway) also had downregulated pTyr levels in afatinib-treated tumors. Although effects of afatinib treatment could be read out with both frozen and FFPE tissues, downregulation of EGFR pathway was more prominent, in terms of fold-change and reproducibility across different tumors, in frozen tissues compared to FFPE counterparts. Indeed, two of the afatinib treated tumors preserved as FFPE even had higher pTyr levels in EGFR and GAB1 compared to their frozen pairs or the vehicle control. Altered signaling in these FFPE tumors was not due to the protein extraction step since frozen tissues processed with FFPE workflow (Frozen-TFE) did not exhibit such strong differences (Figure S3).

**Figure 3.**
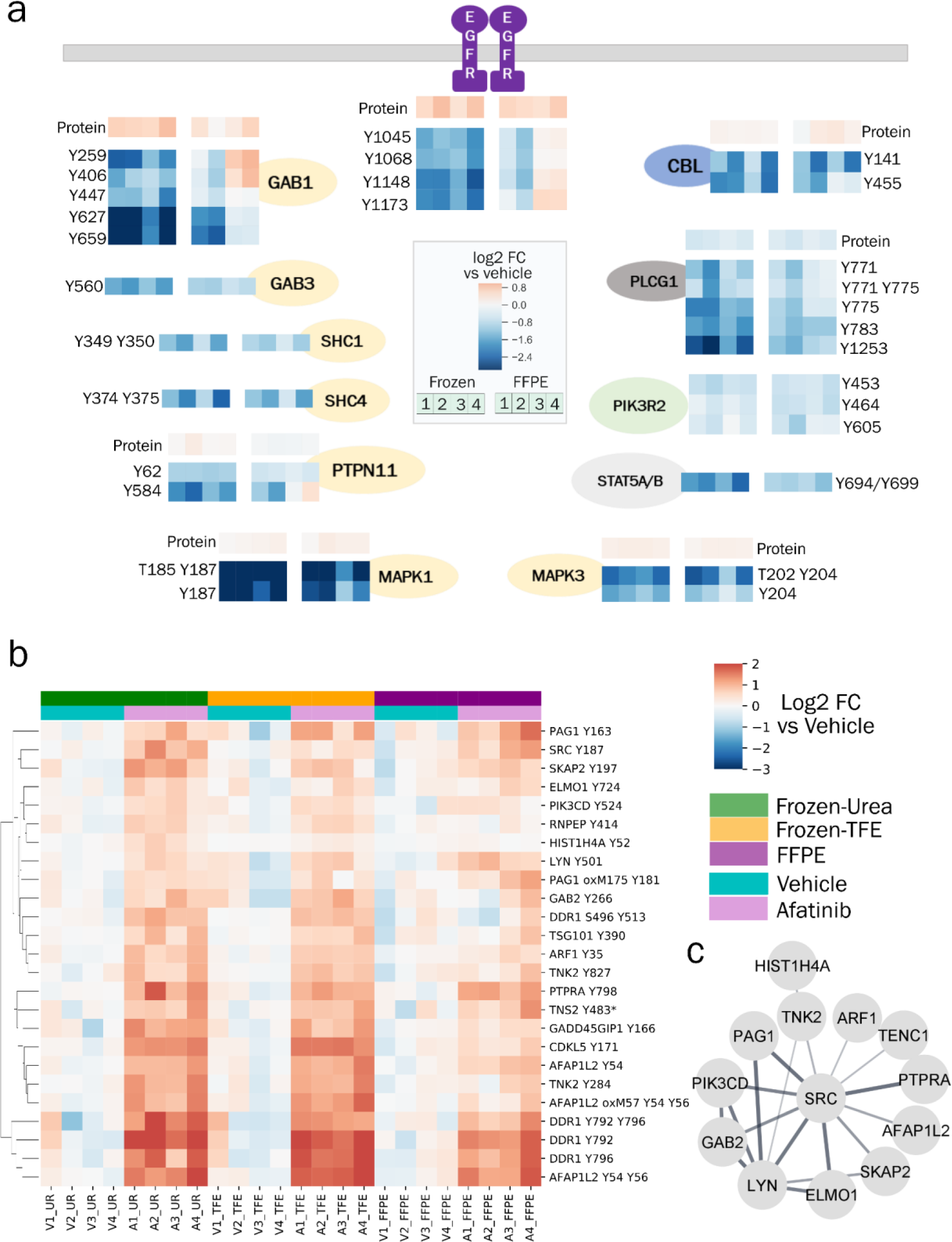
Changes in pTyr and protein levels in response to afatinib treatment. **a)** Diagram of EGFR pathway showing the effect of afatinib treatment on selected pTyr sites in various proteins as quantified in Frozen-Urea and their FFPE counterparts. Protein or pTyr levels are represented as log2 fold change relative to vehicle control. **b)** Hierarchical clustering heatmap of pTyr sites that were significantly upregulated (fold change > 1.4 and FDR q-value < 0.05 after Benjamini Hochberg multiple hypothesis testing correction) in response to afatinib treatment in Frozen-Urea workflow. Phosphotyrosine levels are represented as log2 fold change relative to vehicle control. Miscleaved peptides are denoted by * next to them. **c)** Interaction network of pTyr-proteins from Figure 3b obtained from STRING database. All of the interactions are at least medium confidence based on all interaction sources except text mining. Non-interacting proteins are not shown.

Although afatinib led to downregulation of pTyr signaling in EGFR pathway, it led to upregulation of EGFR and GAB1 at protein level (Figure 3a), highlighting a potential therapeutic resistance pathway where tumor cells upregulate the protein target to overcome loss of activation. Additionally, in response to afatinib, tumors exhibited higher pTyr levels in epithelial discoidin domain containing receptor 1 (DDR1), the non-receptor protein tyrosine kinase Src, and multiple Src-family kinase substrates (Figure 3b), in agreement with previous work demonstrating Src and Src-family kinases (SFK) as a resistance mechanism for EGFR inhibition^21,23^. Phosphoproteins with increased pTyr levels in response to afatinib formed a strong interaction network associated with RTK signaling and locomotion pathways (Figure 3c). These adaptive resistance pathways could be read out from frozen tissues and their FFPE counterparts, suggesting that, despite some signaling alterations during preservation, pTyr analysis of FFPE tissues can provide similar biological information compared to their frozen counterparts.

### Phosphotyrosine signaling is preserved in clinical specimens

Although the data from freshly prepared FFPE and frozen PDX tumors suggested the potential capability of extracting activated signaling networks from pTyr analysis of FFPE tissues, we wanted to assess whether we could obtain similar data from archived FFPE tissues. We therefore expanded the comparison analysis to clinical specimens derived from the Mayo Clinic Breast Cancer SPORE tissue registry where matched frozen and FFPE specimens are collected prospectively, in order to evaluate whether pTyr signaling in FFPE tissues is similar to concomitantly obtained frozen tissues. We obtained 20 triple negative breast cancer (TNBC) tissue specimens, including 10 tumor samples wherein FFPE tissue blocks and matched flash-frozen tumor samples were harvested in parallel from the same tumors (Figure 4a). Two-millimeter punches of tumor-rich regions from the FFPE tissue blocks were obtained by pathologists. Tumor-rich content was verified in the flash-frozen OCT-embedded tissues by pathologist evaluation of haematoxylin and eosin-stained cryosections. Proteins were extracted from each tissue type with the corresponding workflows, digested to peptides, and labeled with isobaric mass tags (TMT10plex). Analysis of enriched pTyr peptides led to identification and quantification of 927 and 382 sites in FFPE and frozen tissues (Table S17-18), respectively, with 281 sites quantified across both sets of analyses. Peptides containing pTyr sites from both analyses clustered by patient (Figure 4b), and the average Pearson correlation coefficient between FFPE and frozen pairs (R = 0.51 ± 0.18) was significantly higher than that of all other pairwise analyses (R = 0.05 ± 0.16, P = 7.56 x 10^−17^) (Figure 4c), indicating that similarity between frozen and FFPE tissues outweighed inter-patient heterogeneity.

**Figure 4.**
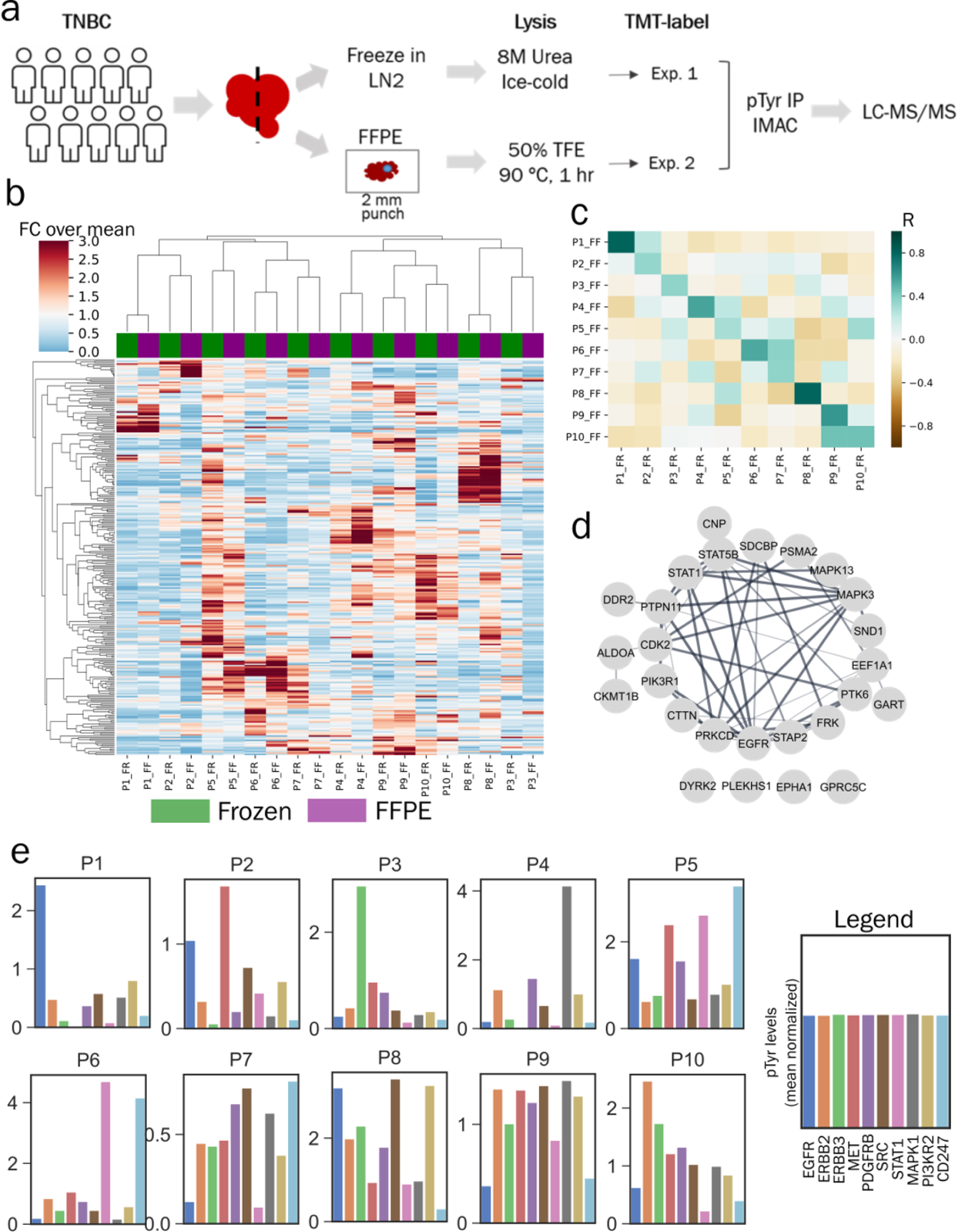
Phosphotyrosine analysis of TNBC clinical specimens. **a)** Experimental workflow to compare pTyr signaling in FFPE and flash frozen specimens from TNBC patient tumors. **b)** Hierarchical clustering heatmap based on Pearson correlation distance metric of pTyr peptides identified and quantified in FFPE and flash frozen conditions. Quantification levels were mean normalized within each workflow before concatenating together. Total of 281 pTyr peptides were quantified in both workflows. **c)** Heatmap of Pearson correlation (R) between flash frozen and FFPE tissues for each patient. Average R for FFPE and frozen pairs (from same patient) was 0.51 ± 0.18 and 0.05 ± 0.16 for all other pairwise analyses. **d)** Interaction network of pTyr-proteins that were highly preserved in FFPE tissues. Phosphotyrosine sites belonging to these proteins had highest Pearson correlation for quantified levels in flash frozen and FFPE specimens. **e)** Barplots with phosphorylation levels of various proteins quantified for each patient based on FFPE tissues. Phosphorylation levels represent average phosphorylation across multiple pTyr sites for a given protein target and are plotted relative to the mean of all 10 tumors (mean normalized).

Despite their relative similarity, the heatmap and correlation coefficients both highlight that FFPE and frozen tumor pairs are not identical. To gain insight into the signaling components that were most highly preserved during FFPE storage, we extracted the sites that had highest correlation between the FFPE and frozen samples. Intriguingly, the top 30 most highly correlated sites belonged to proteins such as EGFR, MAPKs, STATs and PI3Ks, many of which are essential nodes in cellular signaling pathways, highlighting that FFPE tissues can still provide crucial information on dysregulated biological networks in the clinic (Figure 4d, Table S19). Least correlated sites belonged to proteins involved in immune regulation and cytoskeletal organization, potentially highlighting dissimilarity between the tumor rich and stromal regions since FFPE tissues were punched from tumor rich regions, while the frozen tissue specimens were not necessarily tumor-enriched given the larger size of tissue (Figure S4).

TNBC patients have poor prognosis and few therapeutic options beyond chemotherapy. Protein targets such as ERBBs, MET, SRC, MAPKs and STATs have been explored as potential therapeutic targets for breast cancer^24^; therefore, we wanted to assess the phosphorylation state of these proteins across the ten FFPE clinical tumor specimens. To quantify phosphorylation, we averaged multiple pTyr sites for each interesting protein target and plotted values relative to the mean of all 10 tumors (Figure 4e). This analysis led to identification of differential activation of proteins in different patients. For instance, patient 1 (P1) had high relative phosphorylation of EGFR, whereas P3 had high relative phosphorylation of ERBB3. P10 had high phosphorylation in both ERBB2 and ERBB3, but not EGFR, suggesting that this tumor may be driven by HER2/HER3 signaling. In contrast, P8 had high relative phosphorylation for all ERBB family members in addition to PDGFRB, SRC and PI3KR2, suggesting multiple potential drivers, or potentially a more heterogeneous tumor. MET was highly upregulated in P2 and P5. This differential phosphorylation of protein targets highlights potential patient specific oncogenic driving kinases, and may indicate the potential benefit of an individualized targeted therapeutic approach for each patient. For instance, an EGFR inhibitor is more likely to have a therapeutic effect in P1, but not in P2, P3, P4, or P10 based on this analysis. We also discovered evidence of T cell immune infiltration and activation in P5 and P6 as assessed by high phosphorylation on the T-cell receptor (CD247), suggesting a potential role of an immune checkpoint inhibitor in these patients. Overall, the comparative pTyr analysis of FFPE and frozen tissues from TNBC patients suggest that FFPE tissues may provide biologically meaningful information similar to frozen tissues in the clinic.

### Analysis of NSCLC FFPE clinical tissue sections

Finally, we performed pTyr analysis on FFPE clinical samples from a lung cancer tumor tissue bank to assess low-input sample feasibility and determine whether pTyr analysis of FFPE lung cancer tumor tissues could reveal information regarding EGFR phosphorylation and activation. We collected FFPE tissues from nine patients with NSCLC that had mutations in EGFR (Figure 5a). Two 10-μm sections were obtained from each tissue specimen, and proteins were extracted and digested to peptides following our FFPE protocol. We obtained peptide yields ranging from 157 to 595 μg from each patient, suggesting that a single 10-μm section would have been enough for most of the patients (Figure S5a). MS/MS analysis of enriched pTyr peptides led to identification of 962 sites (Table S20), including 70 kinases spanning across multiple kinase families, including Erbbs (EGFR, ERBB2, ERBB3), focal adhesion kinases (PTK2, PTK2B, PTK6) and MAPKs (MAPK 1, 3, 7, 9, 11-14) (Figure S5b). Since tumor tissues were collected from patients with EGFR mutations, we first quantified pTyr levels on EGFR to examine any correlation between the genomic mutations and phosphorylation / activation of EGFR. Genomic mutations were not correlated with EGFR phosphorylation. For instance, P4, P7, and P9 all had L858R mutation; however, P7 and P9 had ∼2-fold and ∼10-fold higher EGFR phosphorylation compared to P4, respectively (Figure 5b).

**Figure 5.**
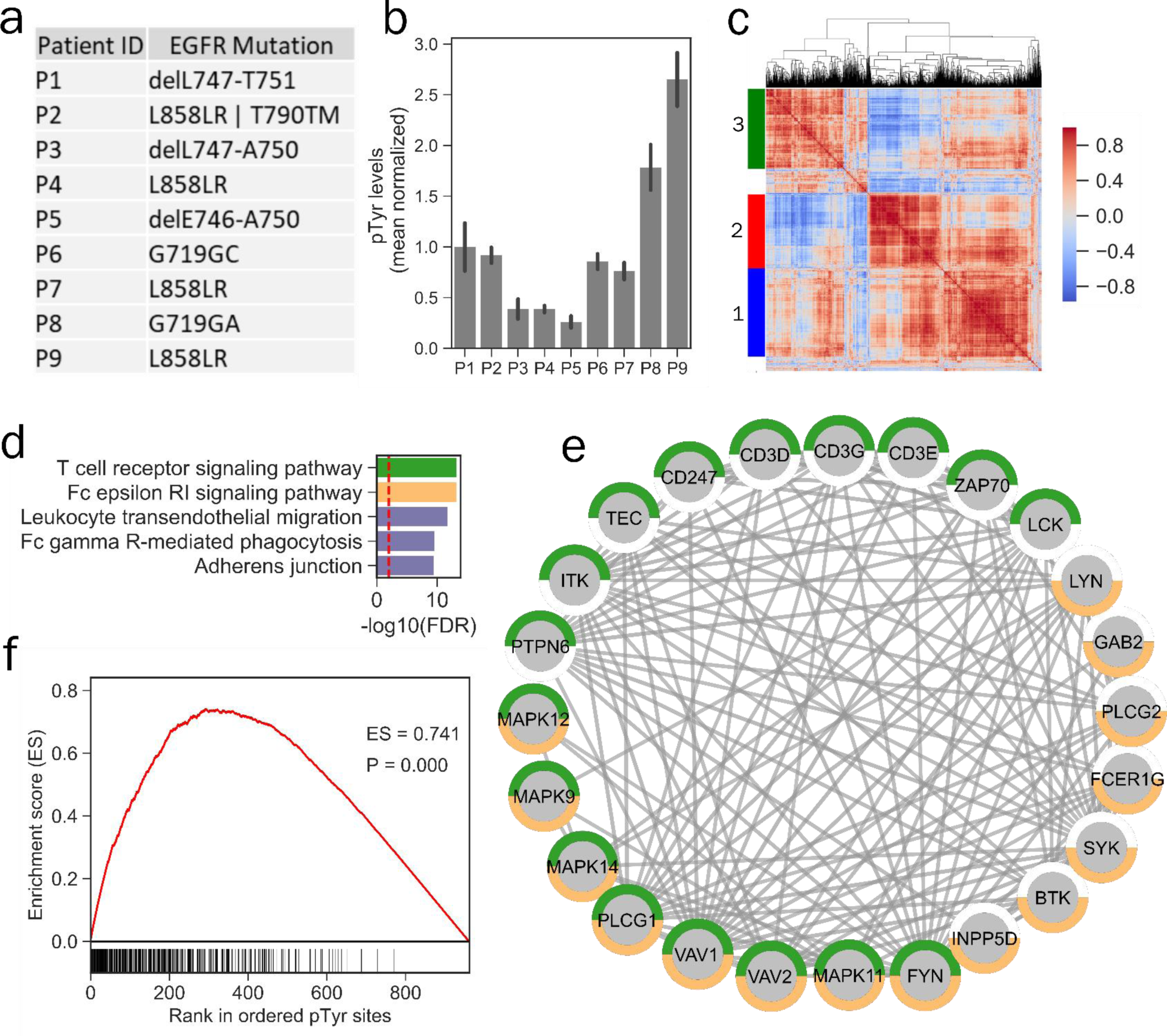
Phosphotyrosine analysis of archived FFPE tissues from NSCLC patients. **a)** Mutation status of EGFR in various patients. **b)** Phosphotyrosine levels of EGFR averaged across multiple tyrosine sites (Y1068, Y1148 and Y1173) plotted relative to the mean of all 9 tumors. Error bars represent standard deviation. **c)** Hierarchical clustering heatmap of co-correlation matrix for pTyr sites quantified in NSCLC FFPE specimens. Clustering was based on Euclidean distance. Color-scale represents Pearson’s correlation. Three main clusters were identified in this analysis. **d)** Top 5 significantly enriched Kegg pathways in pTyr-proteins belonging to cluster 1. Dashed red line depicts FDR q-value = 0.01. **e)** Interaction network of proteins belonging to T cell receptor (green) and Fc epsilon RI (orange) signaling pathways that were identified in cluster 1. All of the interactions are highest confidence based on all interaction sources except text mining from STRING database. **f)** Cluster set enrichment analysis for cluster 1 pTyr sites in Patient 9 (P9).

To identify activated signaling networks in each tumor and extract co-regulated sites, we performed hierarchical clustering on the co-correlation matrix for individual pTyr sites. This co-correlation and clustering analysis revealed 3 main clusters (Figure 5c, Table S20): cluster 1 was highly enriched in innate and adaptive immune signaling, including Fc epsilon RI and T cell receptor signaling pathway consisting of a well-characterized network including proteins such as T cell receptor (CD247, CD3D, CD3E, CD3G), ZAP70 and LCK (Figure 5d-e), whereas cluster 2 was mainly enriched in focal adhesions and regulation of actin cytoskeleton pathways consisting of proteins such as RHOA, ROCK2 and ITGB1 (Figure S5c-d), and phosphoproteins in cluster 3 consisted of several ribosomal proteins as well splicing factors (Figure S5e-f). In order to identify patients with these differentially activated signaling networks, we performed enrichment analysis^25^ for each cluster in each patient. Cluster 1 was enriched in P6, P8 and P9, suggesting immune infiltration and activation in these tumors, which also featured high phosphorylation levels of many other RTKs (Figure 5f, S6a-b). Interestingly, cluster 2 and cluster 3 were highly enriched in single patients, P1 and P2, respectively, suggesting that these tumors may be driven, in part, by splicing and ribosomal dysregulation and integrin / focal adhesion signaling (Figure S6c-d). Together, these data suggest that EGFR mutation status may be a poor predictor of EGFR phosphorylation and activation, as has been suggested previously^26^, and also highlight the potential for direct translational insight from pTyr analysis of 1-2 10-μm sections from FFPE tumor tissue specimens.

## Discussion

FFPE tissues are widely available in the clinic and represent a rich resource to study molecular mechanisms of various diseases directly in patient specimens. Although MS-based proteomics methods have been increasingly used for FFPE samples, these studies have been limited to protein expression and global phosphorylation profiling, both of which are present in high abundance in cells. Identification and quantitation of low-abundant post translational modifications such as pTyr requires additional enrichment and increased sensitivity. We developed a technique to quantify pTyr regulated pathways to provide insight on tumor biology and inform on driving kinases. To the best of our knowledge, this is the first study to quantify tyrosine phosphorylation on several hundred proteins using 1-2 10-μm sections of FFPE. In developing this method, the use of SP3 beads for sample processing and digestion provided a ∼20-fold increase in sensitivity relative to previous publications^27–31^, enabling quantification of pTyr peptides from 50 micrograms of peptides from a single 10-μm FFPE section. As pTyr signaling regulates many aspects of cell and tumor biology, the majority of the identified phosphoproteins belong to several well-characterized cancer pathways, highlighting that pTyr analysis can identify known actionable therapeutic targets from FFPE tissues.

The effects of FFPE preservation on pTyr signaling have not been previously characterized. By comparing the pTyr profiling in FFPE tissues and flash frozen tissues derived from same tumor, we show that FFPE tissues can provide biologically relevant data compared to their flash frozen pairs. While pTyr signaling was more affected by formalin fixing time relative to pSer/Thr or protein expression, the effects of afatinib treatment could still be determined from FFPE tissues. Comparison of pTyr levels between archived flash frozen and FFPE tissues from TNBC patients further substantiated our findings from PDX analyses and highlighted that clinical FFPE tissues could provide similar information compared to their frozen counterparts. However, some caution has to be taken while interpreting the data from FFPE tissues alone since some changes associated with formalin fixation may alter physiological signaling. Additionally, delays in freezing or formalin fixing after surgical resection can affect phosphorylation-mediated cell signaling networks, and thus uniform sample collection procedures are required in the clinic.

Formalin fixing and paraffin embedding is a universal technique for tissue preservation following tissue biopsy and surgical resection, and thus FFPE tissues are readily available. Exome sequencing is commonly used to identify genetic alterations, and transcript profiling typically serves as a proxy for pathway activation. Here we highlight the capability of pTyr analysis of FFPE tissue sections as a method to directly measure signaling network activation in tumors. This approach is amenable to retrospective analysis of clinical specimens, and may be useful to highlight signaling networks associated with therapeutic response/resistance, or to freshly acquired tissues in the clinic for both patient stratification as well as assessing early response to therapeutic interventions.

## Methods

### Animal studies

Studies involving animals were approved by the Institutional Animal Care and Use Committee at Mayo Clinic. The GBM6 PDX cell line was maintained in short-term explant cultures in FBS containing media prior to injection. The GBM6 model (Mayo Clinic PDX National Resource) was injected into the flank of athymic nude mice at a density of 2 million cells per animal (1:1 ratio of cells and Matrigel). Once tumors reached 200-250 mm^3^ (∼14 days), animals were treated with 24 mg/kg of afatinib or vehicle control by oral gavage once daily for 3 days. Each animal received 3 doses, and tumors were harvested 2 hours after the last dose. Half of the resected tumors were flash frozen in LN2 and stored at −80 °C, while the other half of the tumor was processed into FFPE blocks. Tumors were fixed in 10% formalin before being exposed to an ethanol gradient and xylene prior to embedding.

### Clinical samples

All human FFPE and flash frozen tissues were obtained in accordance with approved protocols. TNBC tissues were collected at Mayo Clinic (with institutional IRB approval) and lung cancer FFPE tissues were collected at Moffitt Cancer Center (#MCC18334).

### FFPE protein extraction and lysis

Thin slices with 10-μm size were sectioned with microtome, and sections were collected in 1.7 mL microcentrifuge tubes. Sections were deparaffinized by washing with 500 μL xylene twice, and then hydrated with 500 μL of ethanol for 5 minutes. The sections were incubated at 90 °C in lysis buffer consisting of 50% 2,2,2-Trifluoroethanol (TFE) in 25 mM ammonium bicarbonate at pH 8.5, 1x HALT Protease and Phosphatase Inhibitor Cocktail (Thermo Scientific) and 10 mM of dithiothreitol (DTT) for 1 hour. Lysates were sonicated for 10 minutes. Thiols were alkylated with 55 mM iodoacetamide (IAA) in dark at room temperature for 1 hour. Proteins were desalted using SP3 beads (Thermo Scientific) as described below.

### Frozen tissue protein extraction

Frozen tumors were homogenized in ice-cold 8M urea supplemented with 1x HALT Protease and Phosphatase Inhibitor Cocktail. Frozen tumors were also lysed with a protocol similar to FFPE protein extraction where frozen tumors were incubated in lysis buffer containing 50% TFE in 25 mM ammonium bicarbonate pH 8.5 and 1x HALT Protease and Phosphatase Inhibitor Cocktail at 90 °C for 1 hour followed by homogenization. Protein concentrations were measured by Bicinchoninic acid assay (BCA, Pierce) according to the manufacturer’s instructions. Disulfide bonds were reduced with 10 mm DTT at 56 °C for 1 hour followed by alkylation with 55 mM IAA for 1 hour at room temperature in the dark.

### Desalting and digestion with SP3 beads

After reduction and alkylation of proteins, lysates were incubated with sera-mag speed beads (SP3) and 50% ethanol for 8 minutes at room temperature. One mg of beads per 10-μm FFPE section and 10 μg of beads per 1 μg of protein from frozen tissues were used. Lysate-bead mix was incubated at magnetic rack for 2 minutes and supernatant was discarded. Beads were washed thrice with 200 μL of 80% ethanol. Proteins were digested for 18-24 hours on beads with sequencing grade trypsin (Promega) in 50 mM HEPES buffer at 1:50 trypsin to protein ratio for frozen tumors and 2 μg trypsin per 10 μm section of FFPE. Peptides were collected in the supernatant by incubating beads on magnetic rack. Peptide concentrations were measured by BCA assay. Peptide aliquots were lyophilized and stored at −80 °C.

### TMT labeling protocol

Peptides were labeled with TMT10plex or TMTpro16plex reagents (Thermo Scientific) in ∼35 mM HEPES and ∼30% acetonitrile at pH 8.5 for 1 hour at room temperature at 1.5:4 peptides to TMT reagents ratio (or higher). Labeling reactions were quenched with 0.3% of hydroxylamine. Samples were pooled, dried in speed-vac and stored at −80 °C.

### Phosphopeptide enrichment

Immunoprecipitation (IP) and IMAC were used sequentially to enrich phosphotyrosine containing peptides. Label free samples were resuspended in IP buffer (100 mM Tris-HCl, 0.3% Nonidet P-40, pH 7.4) with 10 mM imidazole. Samples were then incubated with TALON metal affinity resin beads (Takara) beads conjugated with 50 μg of Src SH2 domain^32^ and 16 μg of Fab derived from 4G10 V312 variant^33^. TMT labeled samples were incubated in IP buffer consisting of 1% Nonidet P-40 with protein G agarose beads conjugated to 24 μg of 4G10 V312 IgG and 6 μg of PT-66 (Sigma) overnight at 4 °C. Peptides were eluted twice, each with 25 μL of 0.2% trifluoroacetic acid (TFA) for 10 minutes at room temperature followed by Fe-NTA spin column based IMAC enrichment.

High-Select Fe-NTA enrichment kit (Pierce) was used according to manufacturer’s instructions with following modifications. Eluted peptides from IP were incubated with Fe-NTA beads containing 25 μL of binding washing buffer for 30 minutes. Peptides were eluted twice with 20 μL of elution buffer into a 1.7 mL microcentrifuge tube. Eluates were concentrated in speed-vac until 1-5 μL of sample remained, and then resuspended in 10 μL of 5% acetonitrile in 0.1% formic acid. Samples were loaded directly onto an in-house packed analytical capillary column (50 μm ID x 10 cm) packed with 5 μm C18 beads (YMC gel, ODS-AQ, AQ12S05).

### LC-MS/MS analysis

Liquid chromatography tandem mass spectrometry (LC-MS/MS) of pTyr peptides were carried out on an Agilent 1260 LC coupled to a Q Exactive HF-X mass spectrometer (Thermo Fisher). Peptides were separated using a 140 min gradient with 70% acetonitrile in 0.2 M acetic acid at flow rate of 0.2 mL/min with approximate split flow at 20 nL/min. The mass spectrometer was operated in data-dependent acquisition with following settings for MS1 scans: m/z range: 350-2000; resolution: 60,000; AGC target: 3×10^6^; maximum injection time (maxIT): 50 ms. The top 15 abundant ions were isolated and fragmented by higher energy collision dissociation (HCD) with following settings: resolution: 60,000; AGC target: 1⨯10^5^; maxIT: 350 ms; isolation width: 0.4 m/z, collisional energy (CE): 33% for TMT labeled and 29% for label free, dynamic exclusion: 20 s. For a global phosphoproteomic and proteomic analysis, half of the supernatants from pTyr IPs were fractionated into 10 fractions as described previously^31^. One-tenth of each fraction was analyzed to quantify protein levels, while rest of the fraction was enriched for phosphopeptides using High-Select Fe-NTA enrichment kit. LC-MS/MS of fractionated samples was performed on an Easy-nLC 1000 coupled to a Q Exactive HF-X mass spectrometer. Peptides were eluted with 80% acetonitrile in 0.1% formic acid using a 90 min gradient. Instrument settings were similar to that of pTyr analysis except top 10 most abundant ions were isolated and fragmented with CE of 29% and maxIT of 150 ms.

Crude peptide analysis was performed on a Q Exactive Plus mass spectrometer to correct for small variation in peptide loadings for each of the TMT channels. Approximately 30 ng of the supernatant from pTyr IP was loaded onto an in-house packed precolumn (100 um ID x 10 cm) packed with 10 μm C18 beads (YMC gel, ODS-A, AA12S11) and analyzed with a 70 min LC gradient. MS1 scans were performed at following settings: m/z range: 350-2000; resolution: 70,000; AGC target: 3×10^6^; maxIT: 50 ms. The top 10 abundant ions were isolated and fragmented with CE of 33% at a resolution of 35,000.

### Peptide identification and quantification

Mass spectra were processed with Proteome Discoverer version 2.2 (Thermo Fisher) and searched against the human SwissProt database using Mascot version 2.4 (Matrix Science). MS/MS spectra were searched with mass tolerance of 10 ppm for precursor ions and 20 mmu for fragment ions. Cysteine carbamidomethylation, TMT-labeled lysine and TMT-labeled peptide N-termini were set as fixed modifications. Oxidation of methionine and phosphorylation of serine, threonine and tyrosine were searched as dynamic modifications. TMT reporter quantification was extracted and isotope corrected in Proteome Discoverer. Peptide spectrum matches (PSMs) were filtered according to following parameters: rank = 1, search engine rank = 1, mascot ion score > 15, isolation interference < 30%, average TMT signal > 1000. Peptides with missing values across any channel for PDXs tumor analysis were filtered out. Phosphorylation sites were localized with ptmRS module^34^ with 216.04 added as a diagnostic mass for pTyr immonium ion^35^. PSMs with >95% localization probability for all phosphorylation sites were included for further analysis. For global proteome analysis, peptides were additionally filtered with FDR (Percolator q-value) < 0.01. Only proteins with either two unique peptides or two PSMs were quantified for downstream proteomic analysis.

### Data analysis

Data analyses were performed in Python (version 3.6) and Microsoft Excel 2016. TMT reporter ion intensities from PSMs were summed for each unique phosphopeptide. For protein level quantification, TMT reporter intensities were summed for all unique peptides. Peptide or protein quantification were normalized with relative median values obtained from crude lysate analysis to adjust for sample loading in TMT channels. Student’s *t*-test was used to determine statistical significance between treatment groups. Unsupervised hierarchical clustering was performed based on Pearson correlation distance metric, unless otherwise specified. Protein networks were obtained from STRING (version 11.0) database^36^ and visualized using the Cytoscape platform (version 3.8)^37^. Gene ontology and Kegg pathway enrichment were performed using STRING and PANTHER (version 15.0)^38^ databases. Kinome trees were obtained from KinMap^39^with illustration reproduced courtesy of Cell Signaling Technology, Inc (www.cellsignal.com). For cluster set enrichment analysis, pTyr sites were rank ordered according to their mean normalized phosphorylation levels compared to all 9 tumors, and running enrichment score (ES) was calculated^25^. Significance (P) of ES was derived from 1000 permutations where ranks of pTyr sites were randomized. P represents fraction of permutations where the maximum ES was greater than the observed one.

## Data availability

The mass spectrometry proteomics data have been deposited to the ProteomeXchange Consortium via the PRIDE^40^ partner repository with the dataset identifier PXD020284 and 10.6019/PXD020284. All other data is available upon request.

## Acknowledgements

We thank members of the White lab for helpful suggestions and the Koch Institute’s Robert A. Swanson (1969) Biotechnology Center for technical support, specifically Histology and Biopolymers & Proteomics core facilities.

## Author contributions

Concept and Design: INK, DMB, JCB, LW, MPG, EBH, JNS, FMW. Method development: INK, JK. Investigation: INK, DMB, ACM, KKB, JMC. Data acquisition: INK. Analysis and interpretation of data: INK, FMW. Writing – original draft: INK, FMW. Writing – review and editing: INK, DMB, ACM, JMC, EBH, MPG, FMW. Supervision: EBH, MPG, JNS, FMW

## Supplementary Materials

**Figure S1.**
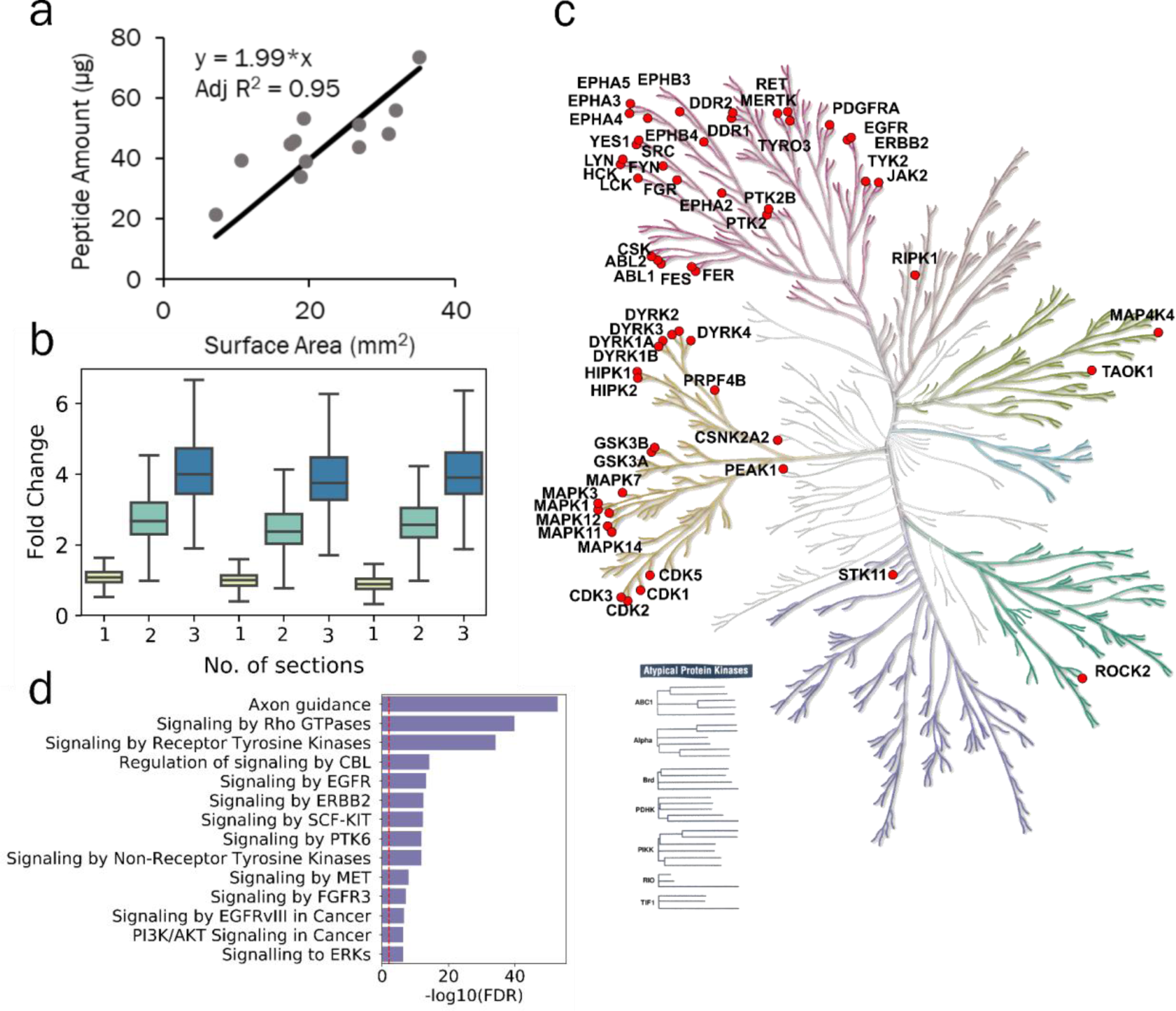
Protein extraction and pTyr analysis in single 10-μm FFPE sections. **a)** Peptide yields from FFPE tissues were proportional to the surface area of tumors. Approximately two micrograms of peptide were derived per mm^2^ of FFPE tissue. **b)** Fold change of TMT intensities of peptides quantified in each channel compared to the average of TMT intensities from single sections from crude lysate analysis. Error bars represent interquartile range. **c)** Kinome tree depicting pTyr containing proteins identified in single 10-μm sections of FFPE tissues. **d)** Selected reactome pathways enriched in gene ontology analysis of pTyr-proteins quantified in single 10-μm FFPE sections. Dashed red line depicts FDR q-value = 0.01.

**Figure S2.**
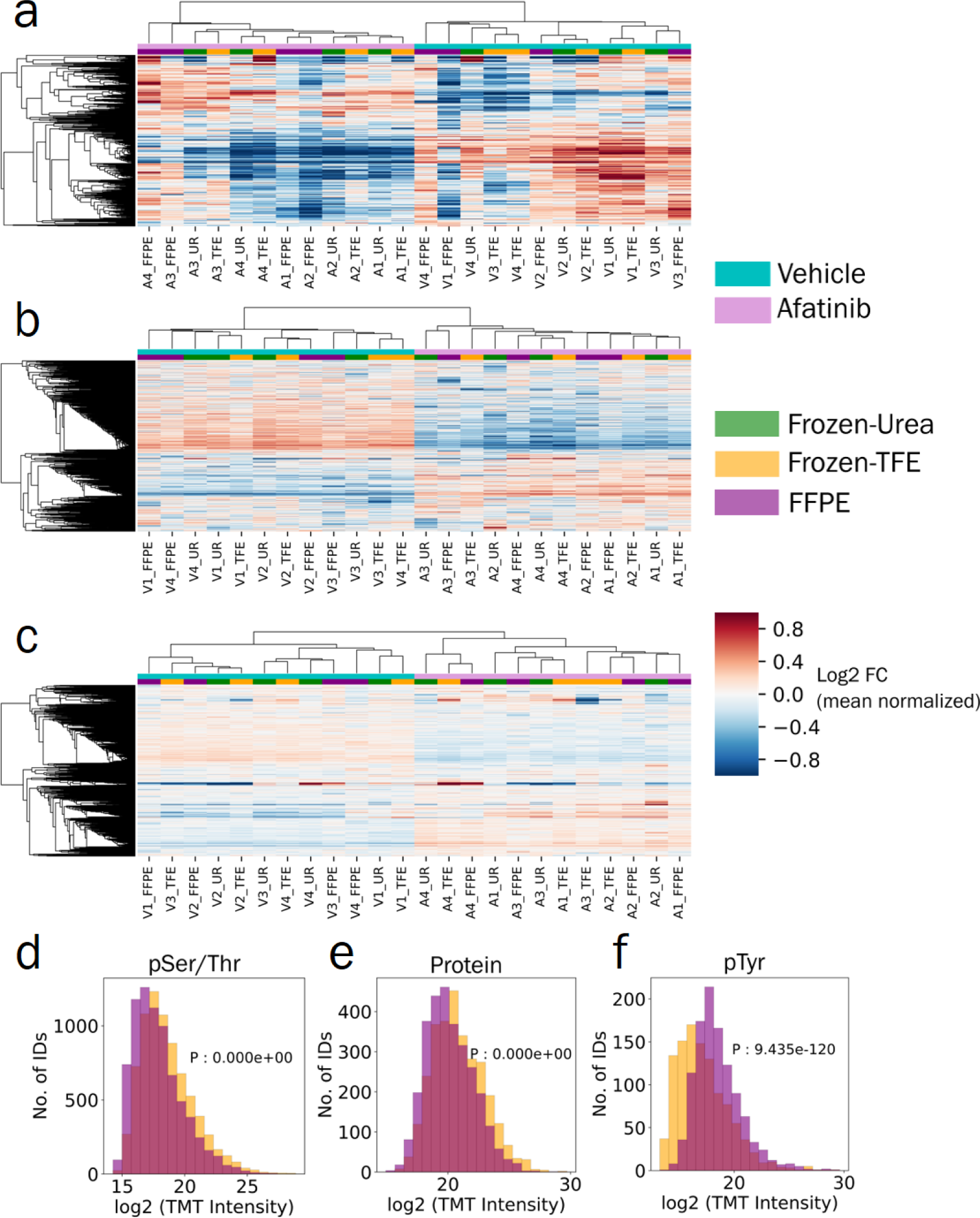
Comparison of phosphoproteomics and proteomics in FFPE and flash frozen tissues. **a-c)** Hierarchical clustering heatmap of **(a)** pTyr, **(b)** pSer/Thr and **(c)** proteins identified and quantified across Frozen-Urea (UR), Frozen-TFE and FFPE workflows. Quantified levels were mean normalized and log2 transformed within each workflow before concatenating together. **d-f)** Differential TMT intensities observed in Frozen-TFE and FFPE workflows across **(d)** pSer/Thr, **(e)** proteins and **(f)** pTyr. TMT intensities were summed across all channels within the workflow. P values were derived from paired two-sided *t*-test.

**Figure S3.**
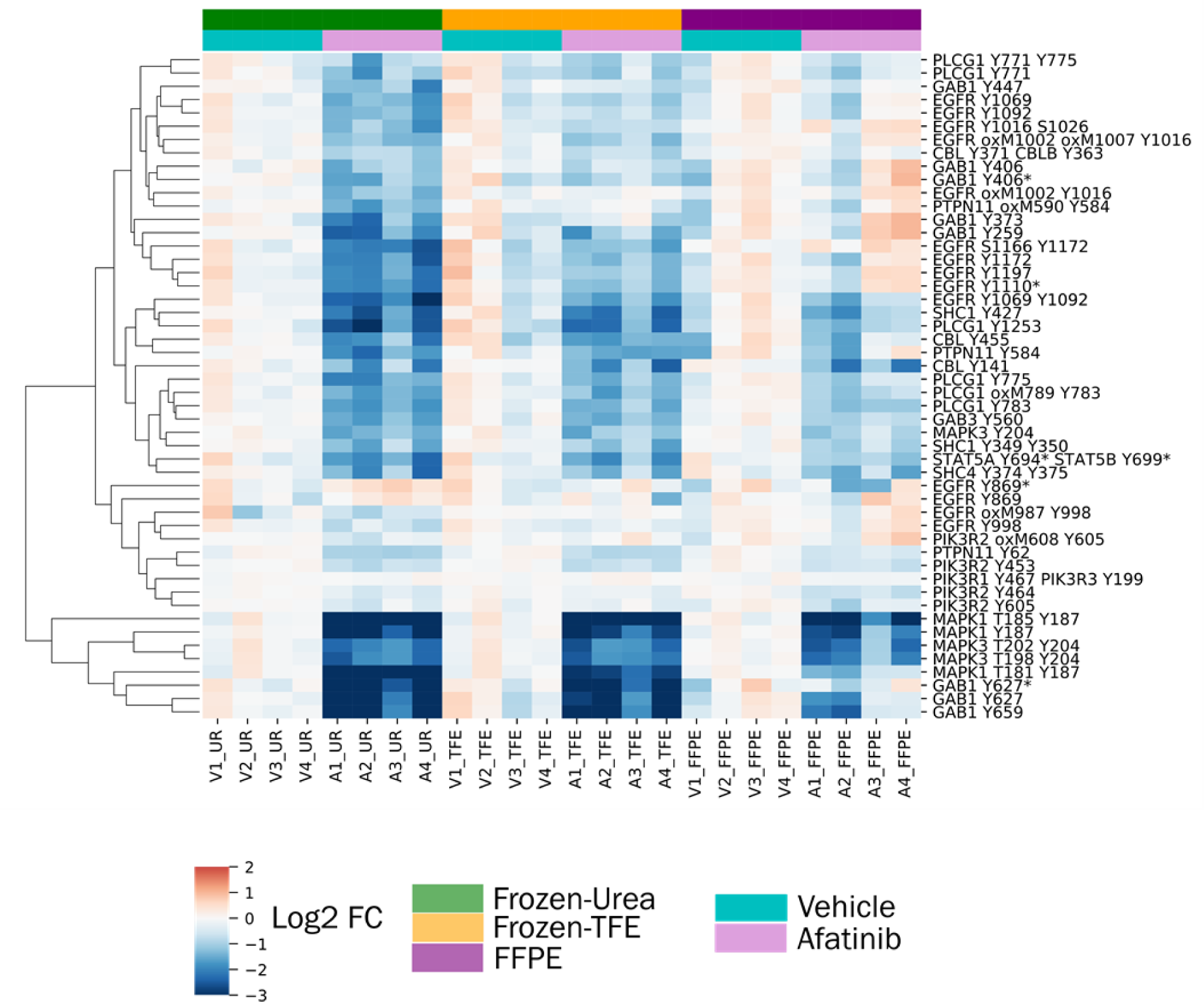
Phosphotyrosine signaling in response to afatinib treatment in selected proteins belonging to EGFR pathway as quantified in Frozen-Urea (UR), Frozen-TFE and FFPE workflows. Quantified levels are presented as log2 fold change relative to the average of vehicle treated group. Oxidation of methionine is denoted by oxM. Miscleaved peptides are denoted by *.

**Figure S4.**
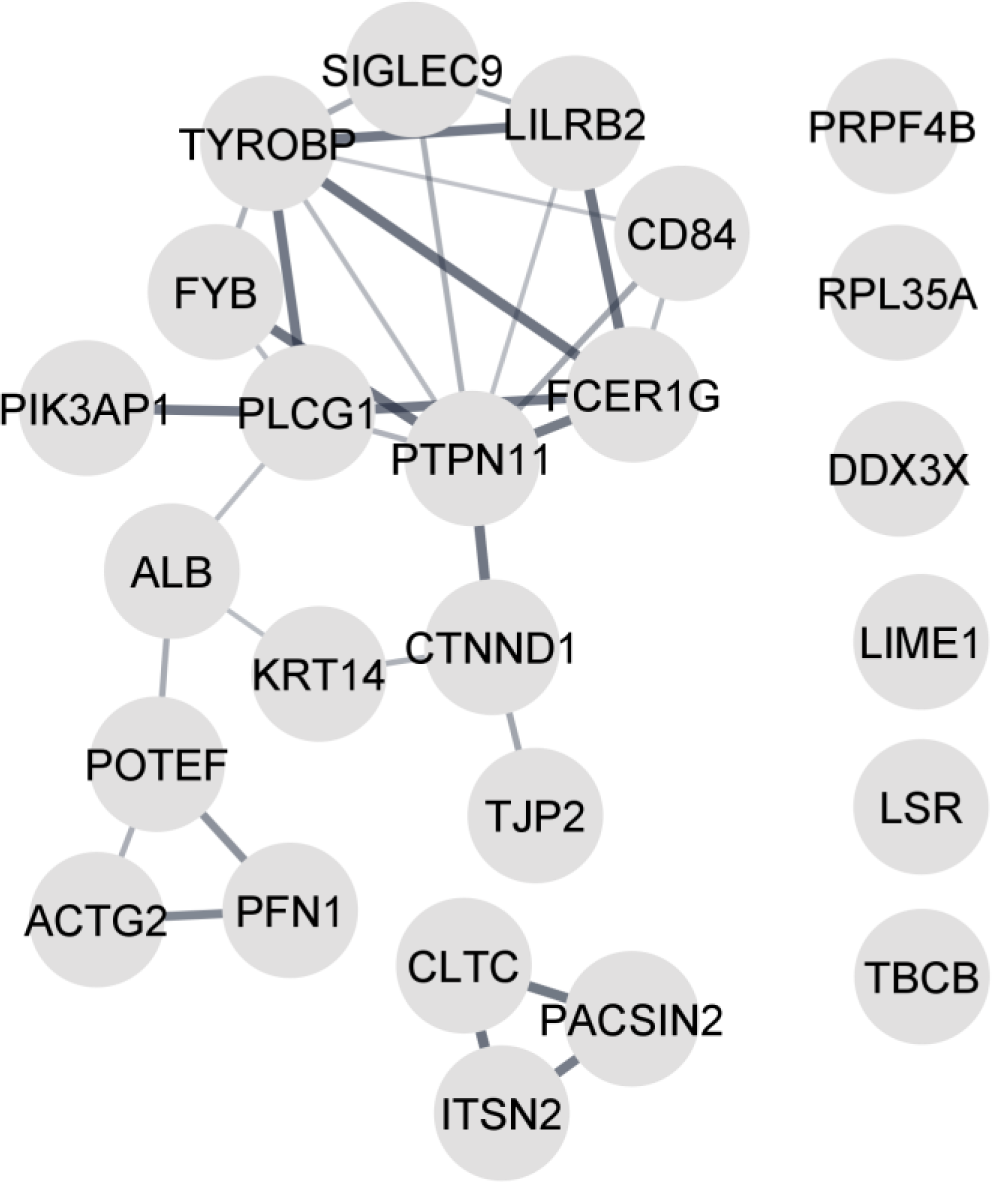
Proteins with pTyr sites that were poorly correlated between flash frozen and FFPE tissues of TNBC patient tumors.

**Figure S5.**
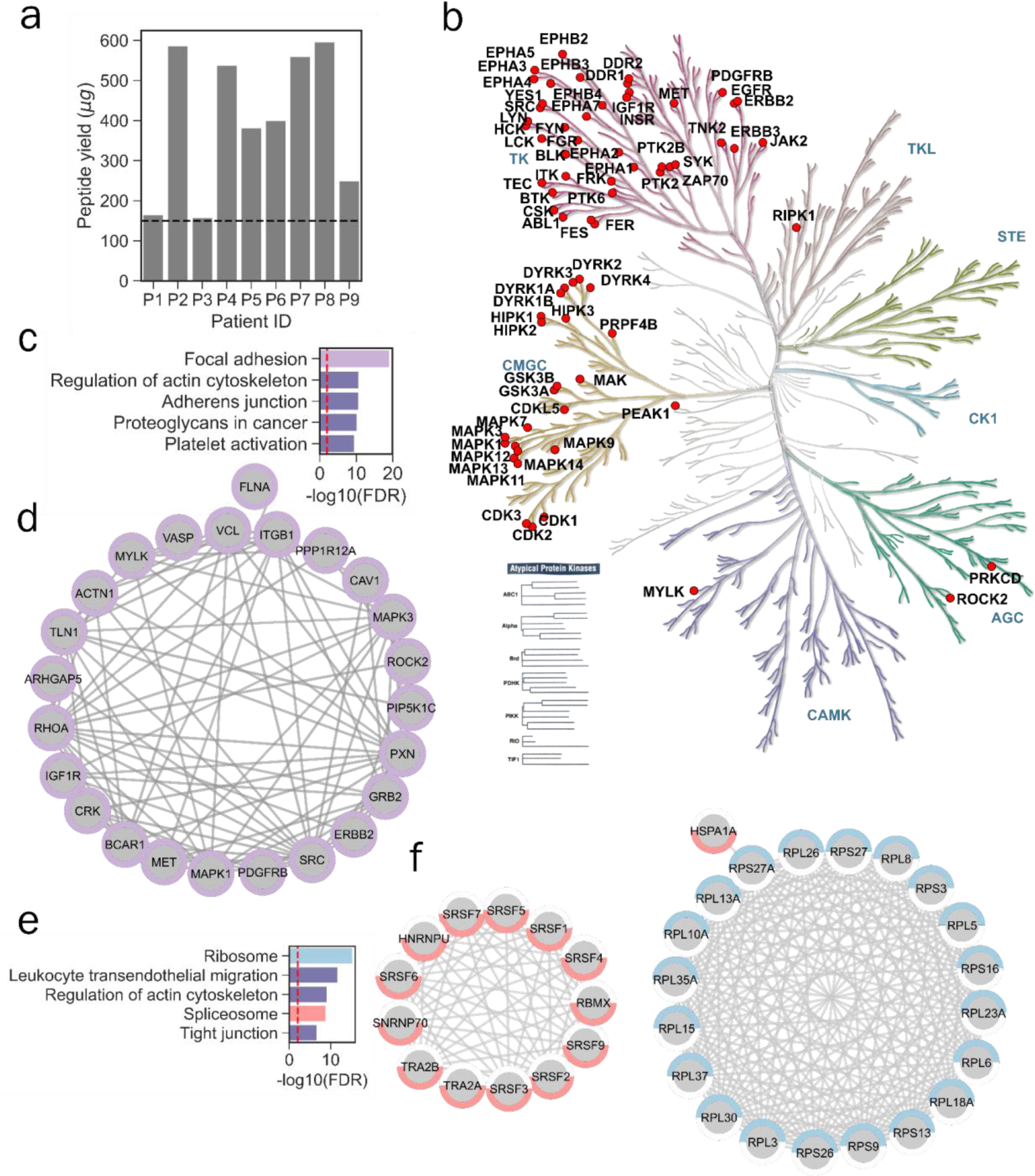
Phosphotyrosine analysis of FFPE specimens from NSCLC patients from a tumor tissue bank. **a)** Peptide yields from 2 10-μm sections of FFPE tissues as measured by BCA assay (average peptide yield = 403 μg). Dashed line depicts peptide amount of 150 μg used for a multiplexed analysis. **b)** Kinome tree depicting pTyr containing proteins quantified in the multiplexed pTyr analysis. **c)** Top 5 significantly enriched Kegg pathways in pTyr-proteins belonging to cluster 2 from Figure 5c. **d)** Interaction network of proteins belonging to Focal adhesion that were identified in cluster 2. **e)** Top 5 significantly enriched Kegg pathways in pTyr-proteins belonging to cluster 3 from Figure 5c. **f)** Interaction network of proteins belonging to Ribosome (cyan) and Spliceosome (red) that were identified in cluster 3. All of the interactions are highest confidence based on all interaction sources except text mining from STRING database. Dashed red line depicts FDR q-value = 0.01.

**Figure S6.**
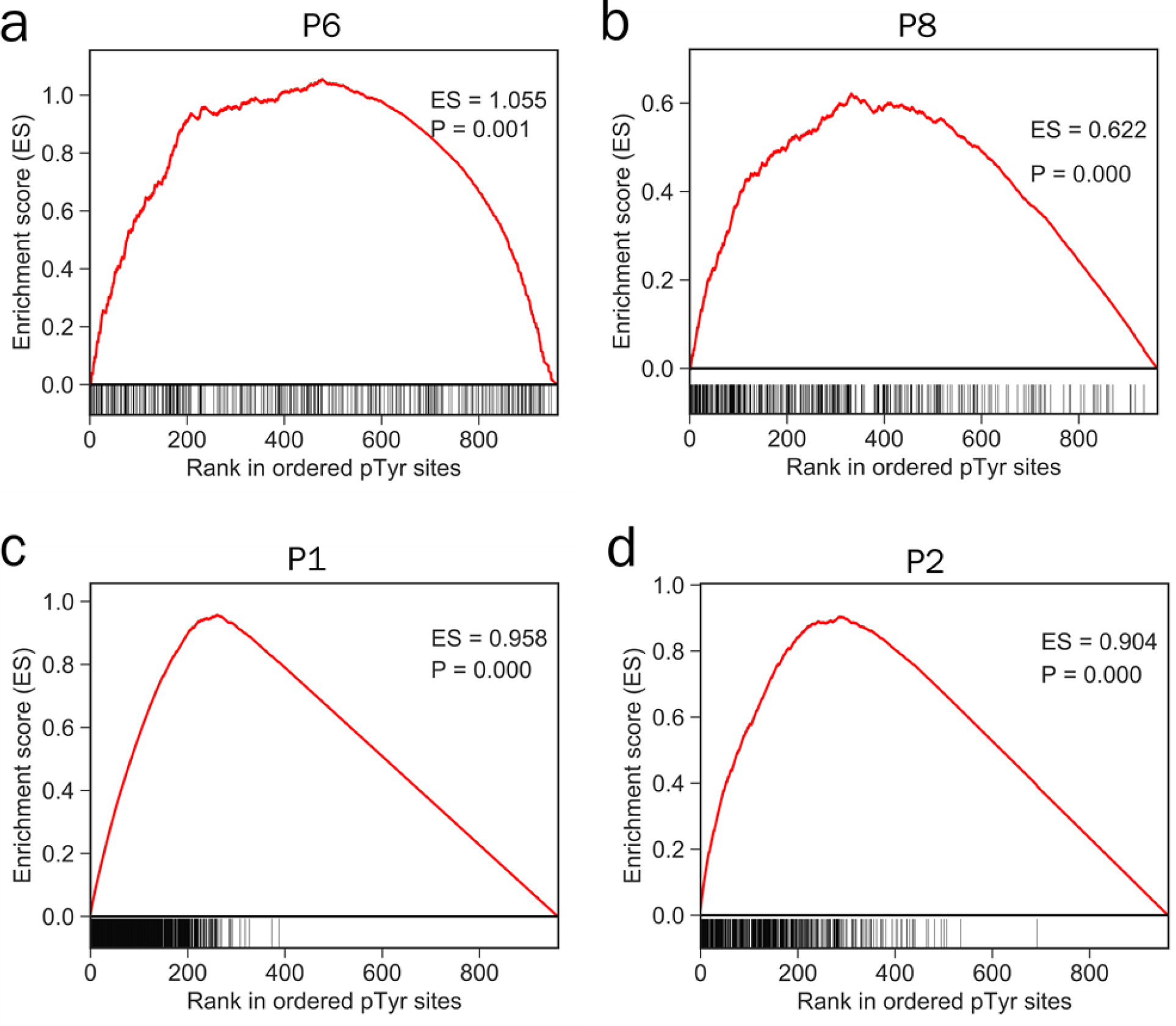
Cluster set enrichment analysis for clusters observed in Figure 5c. Enrichment of cluster 1 in **(a)** P6 and **(b)** P8. Enrichment of **(c)** cluster 2 in P1 and **(d)** cluster 3 in P2. Phosphotyrosine sites were rank ordered according to their mean normalized phosphorylation levels compared to all 9 tumors, and running enrichment score was calculated. Significance (P) of ES was derived from 1000 permutations where ranks of pTyr sites were randomized. P represents fraction of permutations where the maximum ES was greater than the observed one.

Data file 1: Tables with quantitative data extracted from proteomics experiments.

